# DNA isolation in bark beetles: reliability of extraction methods and application in downstream molecular procedures

**DOI:** 10.1101/2024.04.06.588396

**Authors:** Leonel Stazione, Victoria Lantschner, Juan Corley, Carolina Soliani

## Abstract

Molecular tools are increasingly used in entomology for several applications such as taxonomy, genetics, ecology and evolution. For these studies, DNA extraction from individual insects is a crucial step, since the yield and quality vary depending on the methods used. Finding an ideal balance between quality and yield is particularly difficult to reach and several constraints must be considered upon selecting the final protocol. Bark beetles (Coleoptera: Curculionidae, Scolytinae) are a diverse group of forest insects for which molecular studies at the individual level are important, given that many species are invasive and may become pests. However, DNA extraction in bark beetles is not easy, given their small size and complex molecules of diverse nature conforming their exoskeleton and present in their digestive tract. Here, we carried out a comparative analysis of DNA extraction performance in five bark beetle species: *Hylurgus ligniperda, Hylastes ater, Orthotomicus erosus, Orthotomicus laricis* and *Cyrtogenius luteus*. By assessing the efficiency of two different protocols, our aim was to establish the best species-specific method for population-level studies. Our results showed that a method’s whole performance mainly depends on the species considered and translates into DNA quantity and quality variance. We also noted that the traditional method showed better PCR efficiency for the smallest species whereas the commercial kit performed better for the larger beetles. Our comparative analysis provides evidence that no single method for DNA isolation is best, and that each particular species requires optimization.

## Introduction

Molecular tools are increasingly used for a suite of applications in the field of entomology including, taxonomy, genetics and evolution, studies of adaptation to abiotic stress, or for tracking invasions (Borowiec et al. 2022; Lagisz et al. 2010; Miller et al. 2009; Roques et al. 2009; Stevens et al. 2019). This type of molecular work often requires DNA extraction from single individuals (Allen 2006; Dittrich-Schroder et al. 2012; Piper et al. 2019) which varies in yield amount and quality according to the extraction methods deployed (Aboul-Maaty & Oraby 2019; Chen et al. 2010).

Traditional extraction methods work efficiently for DNA isolation from animal tissues but use toxic chemicals and are time-consuming (Dairawan & Shetty 2020; Hajibabaei et al. 2005). In turn, commercially available DNA extraction kits are becoming increasingly popular because of their ease-of-use for a wide range of biological samples, limited labor, and ability to consistently produce high-quality DNA (Nishiguchi et al. 2002; Hajibabaei et al. 2005). However, some of the drawbacks of these kits include their expense and the fact that prolonged incubation times cannot be avoided in any case (Ball & Armstrong 2008). Therefore, an ideal balance between quality and yield is desirable and, at the same time, methodological constraints should be considered upon selecting the final protocol (e.g., the starting tissue, or the size of specimens; Chen et al. 2010; Dittrich-Schroder et al. 2012; van der Loos & Nijland 2021).

Bark beetles (Coleoptera, Curculionidae, Scolytinae) are a highly diverse group of forest insects that breed and feed on the phloem of trees (Kirkendall et al. 2015; Gomez et al. 2023). These insects are characterized by a small body size -most species range from 1 to 5 mm long-. Several species within this group are widely acknowledged as significant agents of tree mortality in coniferous forests globally (Raffa et al. 2008; Biedermann et al. 2019). Additionally, they are inadvertently transported via wood products and wood packaging materials, and numerous species have effectively established worldwide, constituting a notable group of invasive forest insects (Lantschner et al. 2020; Grégoire et al. 2023).

Every extraction method has a percentage of DNA lost during procedure that should be accounted for, to obtain a minimum amount of DNA for analysis. In the case of bark beetles, the main constraints are related to their size and the presence of undesired macromolecules. Their small size imposes a limitation to physical breakdown (milling) that then translates into a lower efficiency of the cell lysis treatment and cell DNA isolation. Working with small amounts of DNA also increases the risk of contamination (van Oorschot et al. 2010). DNA contaminants can be introduced during sample processing, consumables manipulation, laboratory reagents preparation (Ishak et al. 2011; Salter et al. 2014; Stinson et al. 2019), or inappropriate use of commercial DNA extraction kits (Gefrides et al. 2010; Goodrich et al. 2014), in a proportion that may exceed DNA, resulting in a useless extract.

Additionally, bark beetles have complex and large molecules of a diverse nature that might be in part isolated with DNA during extraction procedures, affecting the amplification process (Ferrara 2023; Hosseini 2010). These molecules are mainly proteins and polysaccharides of their exoskeleton -as in most other insect species-, as well as plant phenolics and tannins present in their digestive tract, associated to their xylophagous diet. Macromolecules of this kind may act as inhibitors of PCR and are therefore undesired (Amini et al. 2021; Demeke & Jenkins 2010). For instance, phenolics, tannins and other contaminants present in a DNA extraction solution can inhibit enzymes such Taq polymerases, making extracted DNA unsuitable for PCR amplification (Asghar et al. 2015). Moreover, the DNA extraction method could be affected by different factors such as the storage conditions of the sample, and the cost and duration of the extraction method (Psifidi et al. 2015).

We carried out a comparative study of two extraction methods for five different bark beetle species: *Hylurgus ligniperda* (Fabricius 1787)*, Hylastes ater* (Paykull 1800)*, Orthotomicus erosus* (Wollaston 1857)*, Orthotomicus laricis* (Fabricius 1792), and *Cyrtogenius luteus* (Blandford 1894). The aim of our work was to understand the relationship between the DNA concentration, yield and purity regarding to PCR efficiency presented by each method, in each species. We provide information to ascertain a suitable species-specific protocol that enables population-level studies.

## Materials and Methods

### Sample collection and preparation

Bark beetles were collected from 17 pine plantation stands of southern South America (Argentina, Chile and Uruguay). Insects were captured by placing traps (i.e., Lindgren funnel traps, Cross vane panel traps) with generic bark beetle lures (i.e., ethanol, α-pinene, and/or turpentine), and by manually collecting from cut pine logs available in some sites (see Lantschner et al., 2024 for further details). All collected samples were conserved in 70 - 95% ethanol and stored at −20°C. A total of 173 individuals of five bark beetle species of different body sizes were obtained: 81 individuals of *Hylurgus* ligniperda (4 to 6 mm), 30 individuals of *Hylastes ater* (3.5 to 4.5 mm), 22 individuals of *Orthotomicus erosus* (3 to 3.8 mm), 20 individuals of *Orthotomicus laricis* (3 to 3.7 mm) and 20 individuals of *Cyrtogenius luteus* (2.2 to 2.4 mm).

### DNA extraction

Before extraction, the individuals were oven dried at 60°C for 15 minutes in sterile 1.5-microcentrifuge tubes to ensure removing ethanol residues. We performed two DNA extraction methods:

<u>Method</u> 1 (M1): This method was based on the protocol of Baruffi et al (1995) with modifications. As a starting point, each sample was pre-washed with 50 μl of extraction buffer (removed afterwards) and then grounded with a new aliquot of 100 μl of extraction buffer (100 Mm NaCL, 200 mM sucrose, 50 mM EDTA, 100mM Tris-HC1 (pH 9.1), 0.5 % SDS) using a glass pestle. After this first step, 200 μl of extraction buffer and 2 μl of proteinase K enzyme (10 mg/ml) were added and incubated at 65 °C overnight. Next, the homogenate was mixed with 56 μl of KAc 8M and incubated on ice for 30 minutes. The preparation was centrifuged at 14000 rpm for 15 min, twice consecutively. The supernatant was recovered to then mix it with a solution of 400 μl chloroform: isoamyl alcohol (24:1), following centrifugation at 14000 rpm for 10 min. This step is crucial to ensure DNA isolation and the proper removal of other non-target molecules. The aqueous phase was transferred to a new microcentrifuge tube and the chloroform: isoamyl alcohol step repeated. The DNA was precipitated with 50 µl of NaAc 3M and 400 µl of 100% ethanol. After centrifugation (14000 rpm for 20 min) the resulting DNA pellet was washed twice with 500 µl 70% ethanol. Finally, the pellet was air-dried and eluted in 50 µl TE (TRIS-EDTA 10:1) buffer.

<u>Method 2</u> (M2): This method was based on the use of a commercial kit DNA Puriprep-T kit (Inbio Highway®, Argentina). The entire body of the insects was grounded with a glass pestle by adding 100 μl of the extraction buffer provided with the kit. Then 180 μl of extraction buffer and 2 μl of proteinase K enzyme (10 mg/ml) were further incorporated and samples were incubated at 56 °C overnight. The day after, the homogenate was mixed with 200 μl of a lysis buffer (provided with the kit) and incubated at 56°C for 15 minutes. This step is essential for proper lysis. Then, an aliquot of 200 μl of 100% ethanol was added to the preparation. The entire volume was transferred to a Colum Micro centrifuge tube and centrifuged at 13000 rpm for 2 min. The DNA is retained in the column whereas the eluted volume was discarded. Column membrane was washed twice with 500 µl of washing buffer (provided with the kit). Finally, DNA was eluted from the membrane with 200 µl elution buffer (10 mM Tris-HCl, 0.5 mM EDTA; pH: 9.0, at 70°C).

### Assessment of extraction success

Extraction methods were evaluated using four variables: DNA concentration, yield, 260/280 ratio, and 260/230 ratio. DNA concentration of all samples was measured on NanoDrop™ 2000/2000c Spectrophotometers (Thermo Fisher Scientific, USA) using 2 μl of DNA extraction solution. Yield was reported as the mean ng DNA/mg tissue by method. Purity of samples were reported as the ratio of the absorbance values at 260 and 280 nm (260/280 are the absorbance wave lengths of DNA and proteins, respectively) and at 260 and 230 nm (260/230, are the absorbance wave lengths of DNA, polysaccharides and other compounds like phenolics, respectively).

### PCR amplification

To assess amplification success, each DNA extraction sample was tested through polymerase chain reaction (PCR). The mitochondrial gene cytochrome oxidase c subunit I (COI) was used as the marker of choice - primer pair from Simon et al (1994) (S1718 fw: 5′-GGAGGATTTGGAAATTGATTAGTTCC-3’; A2411 rev: 5′-GCTAATCATCTAAAAACTTTAATTCCWGTWG-3′). The PCR reaction consisted of 1 × buffer (50 mm KCl, 10 mm Tris–HCl, pH 9), 1.5 Mm of MgCl2, 0.2 Mm of each dNTP, 0.16 µM of each primer, 0.6 mg/ml of bovine serum albumin (BSA), 0.04 U of Taq polymerase (Invitrogen®) and 20 ng of DNA, in a final volume of 15 μL. The temperature cycles for the amplification were as follows: initial denaturation at 94 °C for 5 minutes; 40 cycles of 94 °C for 1 minute, 58 °C for 45 seconds, 72 °C 45 seconds and a final extension of 10 minutes at 72 °C. The PCR products (∼ 700 bp) were loaded into 1.5 % agarose containing GelRed Nucleic Acid Gel Stain 10,000X (Biotium®) and electrophoresed in 0.5 × TBE buffer for 1 h at 3 V/cm, together with standard DNA ladders (1 Kb plus; Promega Corp.) to confirm amplification. Samples were purified by using a precipitation method based on sodium acetate solution and then sequenced with an ABIPRISM 3700 capillary sequencer (Applied Biosystem).

### Data analysis

Differences in the extraction success variables (DNA concentration, yield and 260/280 and 260/230 purity ratios) between both extraction methods (M1 and M2) were tested for each species individually. For that, a generalized linear model (GLM) was performed using the logit link function and a Gaussian distribution (best fitted distribution of the data). The extraction method was included as a fixed factor, and the replicate samples - within the method-as a random factor. In addition, a Spearman correlation analysis was carried out to explore the relationship between DNA concentration, yield and purity ratios. Finally, for each species, the proportion of samples for which successful amplifications were obtained with each extraction method was estimated and a Chi-Square (*X*^2^) Goodness of Fit Test was performed to test for differences between methods. All analyzes were performed using R software (version 4.2.2) for Windows (R Core Team, 2023).

## Results

The results of the GLM showed that the success of the two extraction methods varied depending on the species (Fig 1). However, it is possible to identify some trending patterns. Significant differences were revealed between the extraction methods for some extraction success variables, with M1 exhibiting a lower average yield and higher 260/280 purity ratio than the M2, for *H. ligniperda* (z = 6.86, p < 0.001 for yield; z = 5.58, p< 0.001 for 260/280), *H. ater* (z = 2.77, p <0.01 for yield; z = 5.22, p < 0.001 for 260/280)*, O. laricis* (z = 5.82, p < 0.001 for yield, z = 6.97, p < 0.001 for 260/280) *and O. erosus* (z = 2.10, p < 0.05 for yield) (Fig 1). DNA concentration showed variable outcomes, with significantly higher values in M1 than M2 for *H. ater* (*z* = 2.44, p < 0.05; and *C. luteus* (z = 2.67, p < 0.01) whereas the opposite occurred for *O. laricis* with M2 showing significant higher values of DNA concentrations than M1 (z = 6.22, p< 0.001; Fig 1). No differences were found in 260/230 purity ratio between methods for any of the five species (Fig 1).

**Figure 1.**
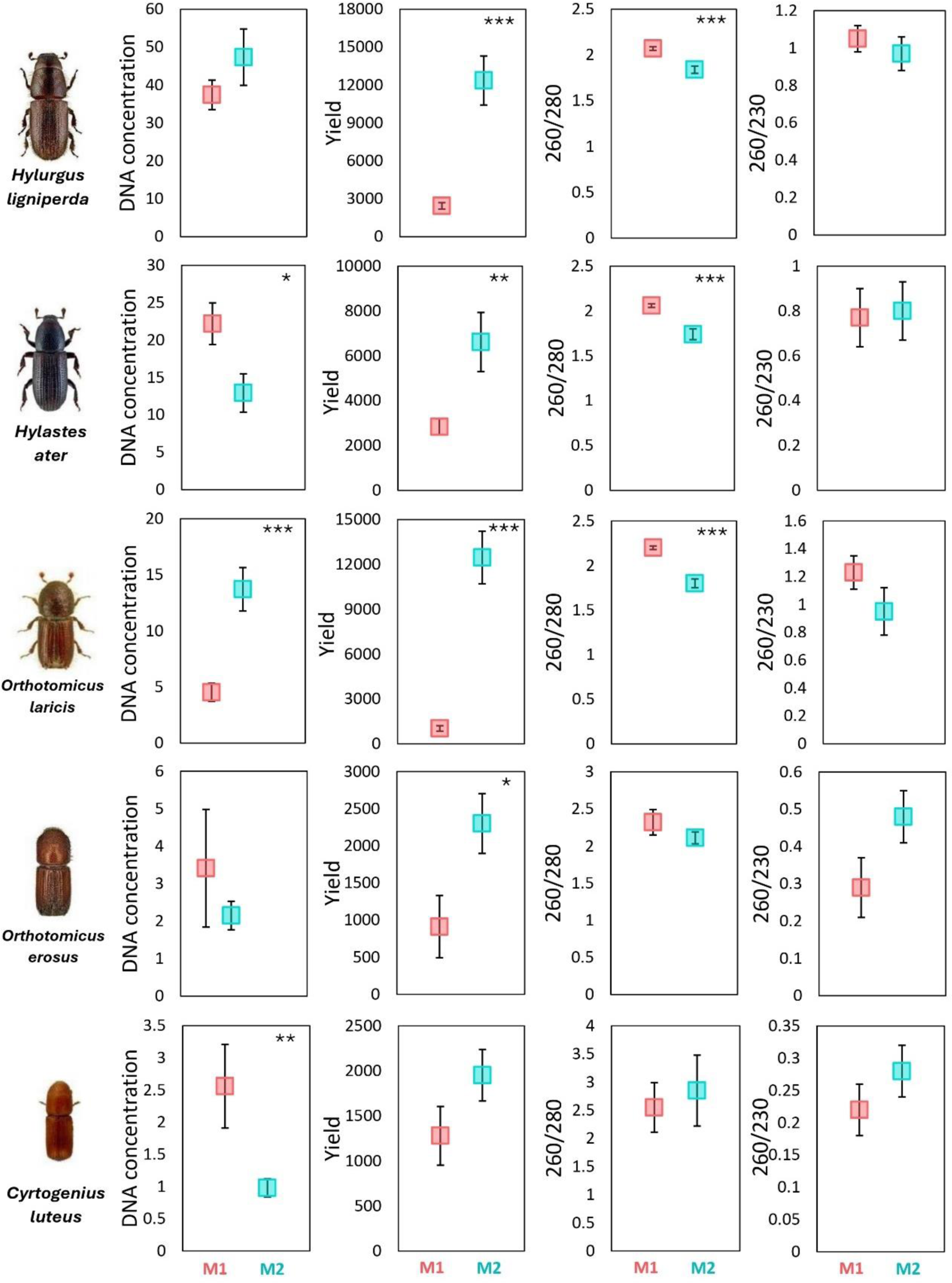
Mean values (±SE) of DNA concentration, yield, and 260/280 and 260/230 purity ratios for each extraction method, in each insect species separately. M1: Baruffi protocol, M2: commercial kit DNA Puriprep-T kit. Asterisks denote significant differences (*p < 0.05, **p < 0.01, ***p < 0.001). The proportionality in the size of the specimens for each species is in accordance with reality

The results of the Spearman correlations between variables of extraction success showed significant positive correlations between DNA concentration and yield in all species. Correlations between DNA concentration and 260/280 purity ratio as well as between yield and 260/280 purity ratio were mostly negative and resulted significant in *H. ligniperda*, *O. laricis* and *C. luteus*. Furthermore, the correlations between DNA concentration and 260/230 purity ratio as well as between yield and 260/230 purity ratio were mostly positive and resulted significant in *H. ligniperda*, *O. erosus* and *C. luteus*. Finally, the correlations between 260/280 and 260/230 purity ratios were significantly positive in *O. laricis* and significantly negative in *C. luteus* (Table 1, Fig S1, S2, S3, S4, and S5).

**Table 1.** Spearman correlation analysis between pairs of extraction success variables: DNA concentration, yield and 260/280 and 260/230 purity ratios. N: number of samples. CC: correlation coefficient. Asterisks indicate significant differences (* p < 0.05; ** p < 0.01; *** p < 0.001).

| Variable 1 | Variable 2 | *H. ligniperda* |  | *H. ater* |  | *O. laricis* |  | *O. erosus* |  | *C. luteus* |  |
| --- | --- | --- | --- | --- | --- | --- | --- | --- | --- | --- | --- |
|  |  | N | CC | N | CC | N | CC | N | CC | N | CC |
| DNA concentration | Yield | 81 | 0.83*** | 30 | 0.55** | 20 | 0.98*** | 22 | 0.79*** | 20 | 0.6** |
| DNA concentration | 260/280 | 81 | -0.36** | 30 | 0.14 | 20 | -0.77*** | 22 | -0.2 | 20 | -0.63** |
| DNA concentration | 260/230 | 81 | 0.37*** | 30 | 0.28 | 20 | -0.21 | 22 | 0.65** | 20 | 0.54* |
| Yield | 260/280 | 81 | -0.65*** | 30 | -0.56** | 20 | -0.81*** | 22 | -0.19 | 20 | -0.58* |
| Yield | 260/230 | 81 | 0.28* | 30 | 0.28 | 20 | -0.27 | 22 | 0.81*** | 20 | 0.9*** |
| 260/280 | 260/230 | 81 | 0.13 | 30 | -0.09 | 20 | 0.45* | 22 | -0.42 | 20 | -0.49* |

A higher proportion of samples showing successful amplifications was obtained for M2 than for M1 in all of species except for *C. luteus*, where the pattern was opposite (*X^2^* = 9.41, p < 0.001 for *H. ligniperda*; *X*^2^ = 86.7, p< 0.01 for *H. ater*; *X*^2^ = 86.7, p < 0.001 for *O. erosus*; *X*^2^ =11.13, p< 0.001 for *O. laricis*; *X*^2^ = 62.5, p < 0.001 for *C. luteus*; Fig 2).

**Figure 2.**
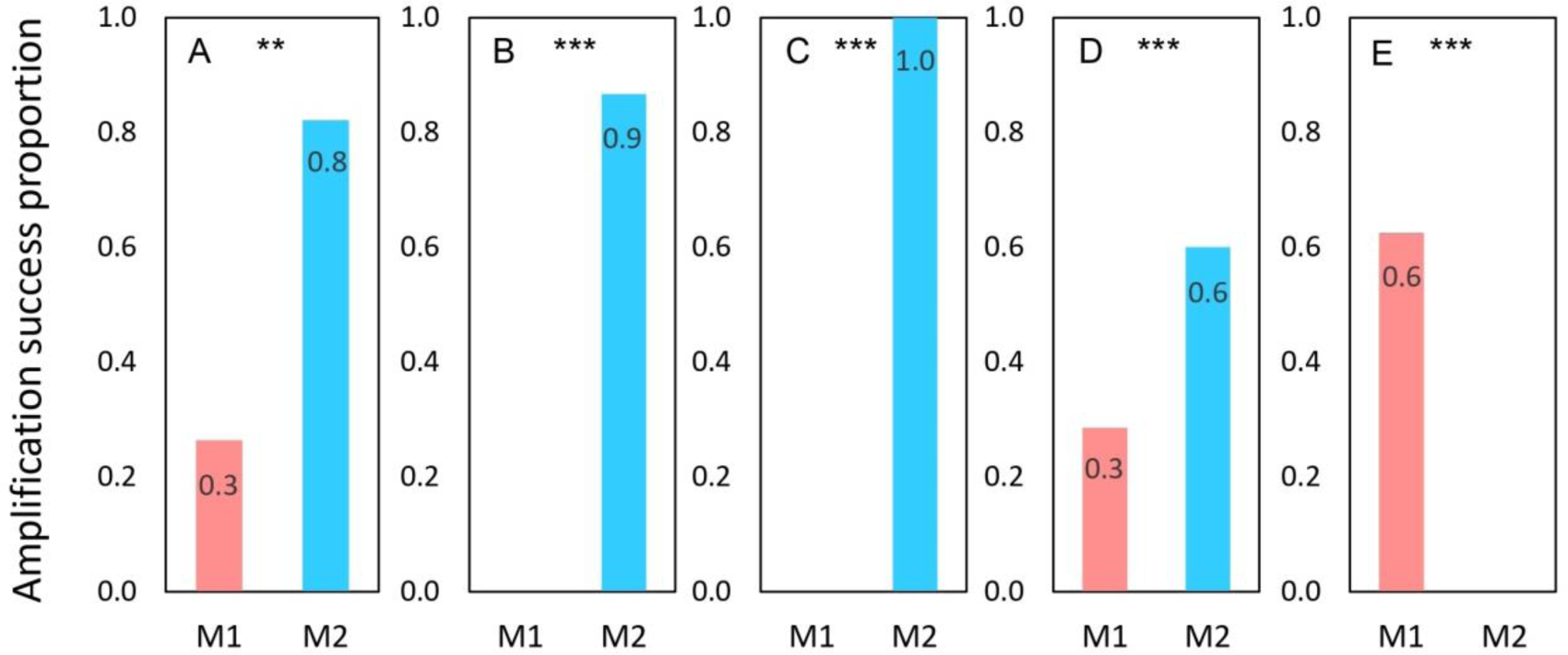
Amplification success proportion for both extraction methods, in each species separately. A: *Hylurgus ligniperda,* B: *Hylastes ater*, C: *Orthotomicus laricis*, D: *Orthotomicus erosus*, E: *Cyrtogenius luteus*. M1: Baruffi protocol, M2: commercial kit DNA Puriprep-T kit. Asterisks denote significant differences (**p < 0.01, ***p < 0.001)

## Discussion

The efficiency of two DNA extraction methods, based on different isolation approximations (phase separation and use of silica membranes) in each of five bark beetle species was different. The DNA quantity and quality values showed a variance that mainly depended on the species under consideration. In general terms, the results indicated that the commercial method performed better in most species. However, in *C. luteus* -the smallest species-the modified Baruffi method showed better PCR efficiency. Therefore, we conclude that it is not possible to recommend a single standard protocol for all species.

We found that DNA concentration is not always predictive of higher PCR performance, and that other properties of DNA quality may be particularly useful when considering the likelihood of amplification success (Ballaré et al. 2019). It has often been assumed that high levels of DNA concentration are necessary for high DNA quality (Andrews et al. 2016; Puritz et al. 2014). However, in this study the method that returned the highest DNA concentration as the most efficient in PCR amplifications was not consistently observed.

When focusing on yield, better results were observed with the commercial kit regardless of the beetle species. It is also important to note that yield is usually influenced by many factors related to species’ characteristics, as for example body sclerotization, preservation method after field collection, or a particular and crucial step during extraction procedure, (i.e., DNA precipitation; Lear et al. 2018). In our case, and although all steps of material conservation, transportation and conditioning were controlled, minor tissue damage could still have occurred that might have influenced the yield and quality of extracted DNA (e.g., Chen et al. 2010), irrespective of the species. These properties of template DNA could become a determining factor for PCR efficiency (Schill 2007). In four out of five species, a higher PCR efficiency was observed in samples that were extracted with the commercial kit. Conversely, in the smallest species, *C. luteus,* a higher PCR success rate was observed for samples treated with M1, in spite of minor yield, opposite to what was observed for the other species.

The 260/280 ratio, a proxy of purity, exhibited a negative relationship with DNA concentration as well as yield in all species, regardless of their size. In addition, the results showed that a higher 260/280 purity ratio did not guarantee PCR efficiency. This trend was consistent across most species, with the method identified as the best-performing candidate (M2) showing a lower PCR success rate but significantly higher 260/280 purity ratio. In this sense, it is important to note that 83.8% of the samples (145 out of 173 total) had a 260/280 purity ratio value higher than 1.8, indicating “pure” DNA with little to no protein or other contaminants that absorb light at or near 280 nm (Simbolo et al. 2013).

In molecular studies that plan to extract DNA from a limited number of individuals, a high PCR efficiency is particularly important. However, in cases where the initial amount of tissue is small and/or the sample quality is low, the extracted DNA may compete with DNA contaminants over the course of an experiment (Lusk 2014). Implementing stricter laboratory precautions, such as routine UV irradiation of reagents and tools and positive pressure laboratory ventilation systems, could help minimize the risk of DNA contamination in the laboratory (Willerslev & Cooper 2005; Champlot et al. 2010). While laboratory contamination is probable, previous studies show that cross-contaminations were most likely introduced from the field (Goldberg et al. 2013; Kranzfelder et al. 2016). The results seem to indicate that in bark beetles the 260/280 purity ratio is not a variable that defines the efficiency of the PCR, but not everything has been said when it comes to tiny specimens that seem exceptional.

The second indicative of DNA purity, the 260/230 ratio, showed a positive relationship with DNA concentration as well as yield in most species. Due to the presence of phenols, polysaccharide and other organic contaminants -that absorb light at 230 nm-the 260/230 ratio could become relevant and tightly link to a particular bark beetle species (Bilgin et al. 2009; Varma et al. 2007). We found opposite trends in purity ratio, probably associated to the performance of each applied method of DNA isolation, between *O. laricis* and *C. luteus*. As the number of PCR successful events were contrasting in both species, we suggest that purity might have determined this outcome relation. Achieving complete elimination of these contaminants is particularly relevant in PCR amplification, given that contaminants cause inhibition in enzymes like restriction endonucleases and Taq polymerases (Arif et al. 2010; Calderón-Cortés et al. 2010; Asghar et al. 2015). Considering the feeding interaction of bark beetles with trees and their frequent exposure to plant products, we propose that there may be a concomitant relationship between DNA concentration and organic contaminant present in the wood during the extraction process. These contaminants might interfere with the efficiency of PCR (Bilgin et al. 2009). Furthermore, this relationship is likely to change depending on the insect size and species. It is expected that the amount of genetic material extracted in smaller species would be less than that obtained in larger species -using the same protocol- and in fact, the proportion of organic contaminants will be greater as the smaller the insect. This could explain why M2 showed higher PCR efficiency in larger species while it failed completely in *C. luteus* – the smallest species-(Fig 4). In addition, the difference between protocols steps could be a key factor determining the PCR efficiency. The Baruffi method includes steps to separate aqueous phases and wash the extracted material with alcohols (see Materials and Methods), which are carefully performed to reduce the losing pellet fractions, especially in small species where the extracted material is even more limited. While in silica membrane-based methods some loss of DNA is reported during the extraction (Rothe & Nagy 2016).

Considering the presence of possible organic wood residues that could interfere with the PCR performance, future tests that include dissection of the digestive tract may be the key to achieve the proper removal of these contaminants. Furthermore, the time elapsed since the DNA extraction was carried out could be an important factor to the PCR performance (Hazir 2019; Soto-Calderón et al. 2009) and should be considered. The DNA isolated from xylophagous insects, which may contain polysaccharides, RNA and phenolics, is not suitable for long storage periods and has a shorter storage life (Horne et al. 2004; Lodhi et al. 1994; Rao & Doddamane 2022).

In bark beetles the presence of residual organic wood molecules and their small and sclerotized bodies represent major constrains in the choice of a reliable extraction protocol. Our results show different key factors to consider when choosing a suitable method in these xylophagous insects. Although there is not a single extraction method for all species, by assessing DNA concentration, yield and purity as estimators of PCR success on several bark beetle species, we provide a novel approach to estimate the applicability of widely used laboratory routines.

## Acknowledgements

The authors thank Silvia Lanzavecchia and the Laboratorio de Insectos de Importancia Agronómica, Instituto de Genética “E.A. Favret”, INTA – CONICET, for their assistance in implementing the Baruffi extraction method (M1); and Sergio Ramos, Edgar Eskiviski, Gonzalo Martínez and Rodrigo Ahumada for providing bark beetle samples. This work was supported by a grant from Agencia Nacional de Promoción Científica y Tecnológica of Argentina (PICT 2019-235) and CONICET (PIP 11220200100764CO).

## Notes

### Competing Interest Statement

The authors have declared no competing interest.

