## Supplemental Figure 1, 2, 3, 4, 5 for "DNA isolation in bark beetles: reliability of extraction methods and application in downstream molecular procedures"

**Supplementary material**

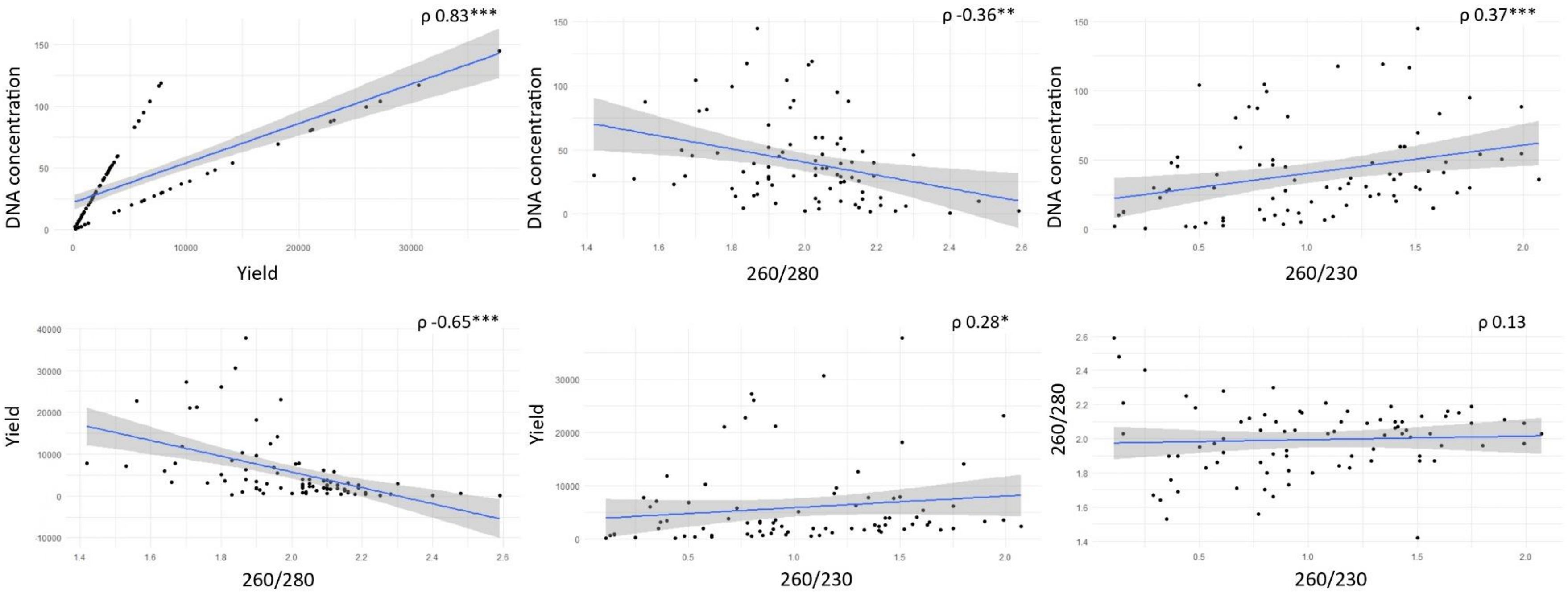


Figure S1. Spearman correlations between all pairs of extraction success variables measured for *Hylurgus ligniperda*. Asterisks denote significant differences (*$p < 0.05$, **$p < 0.01$, ***$p < 0.001$)

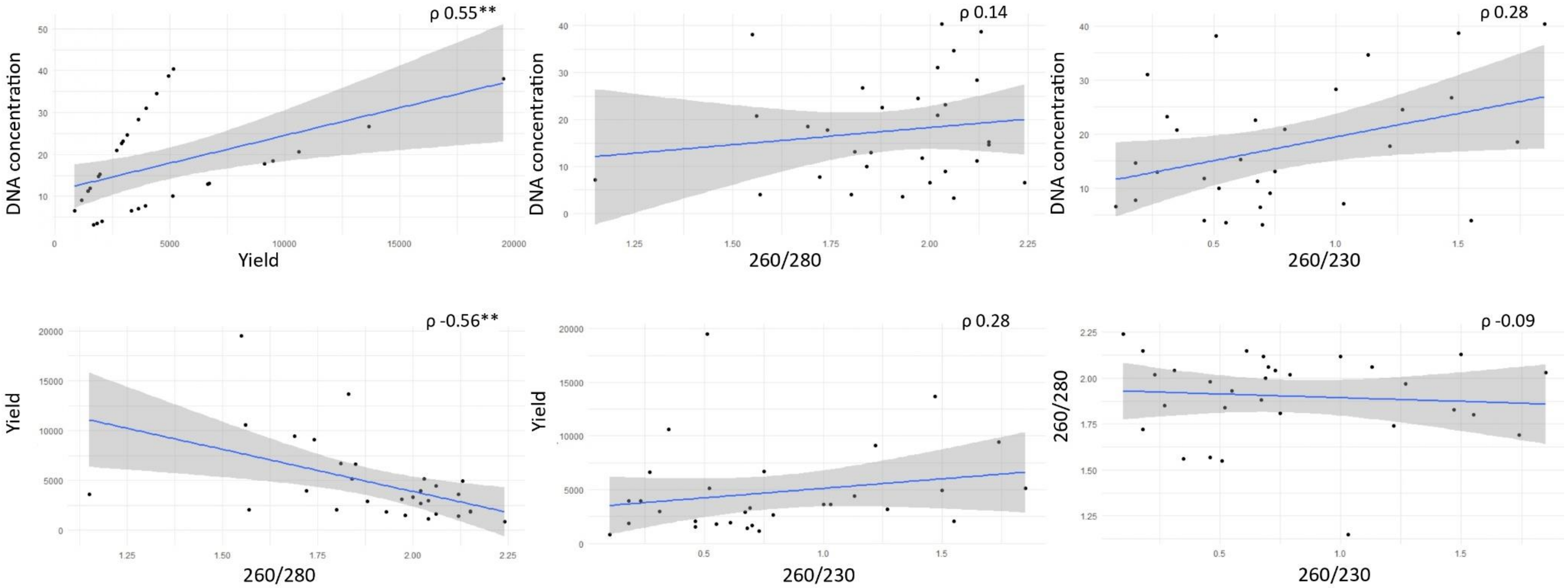


Figure S2. Spearman correlations between all pairs of extraction success variables measured for *Hylastes ater*. Asterisks denote significant differences (*p < 0.05, **p < 0.01, ***p < 0.001)

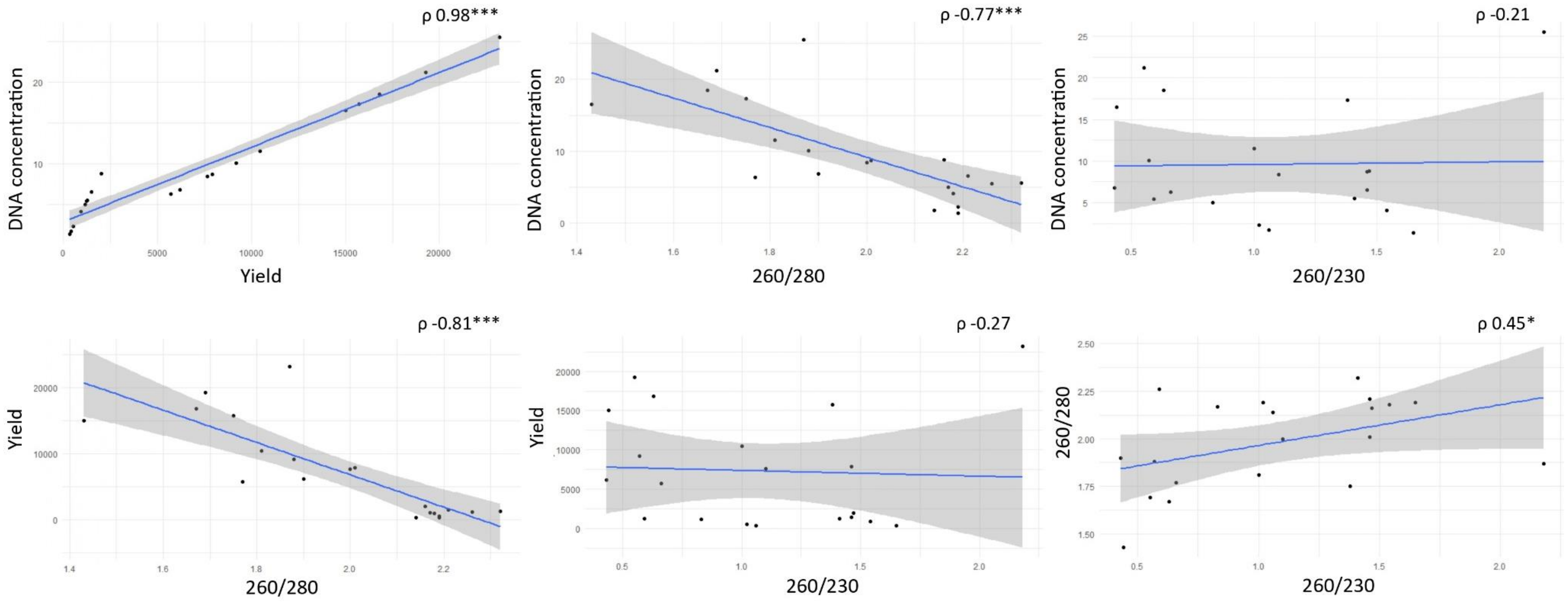


Figure S3. Spearman correlations between all pairs of extraction success variables measured for *Orthotomicus laricis*. Asterisks denote significant differences (*p < 0.05, **p < 0.01, ***p < 0.001)

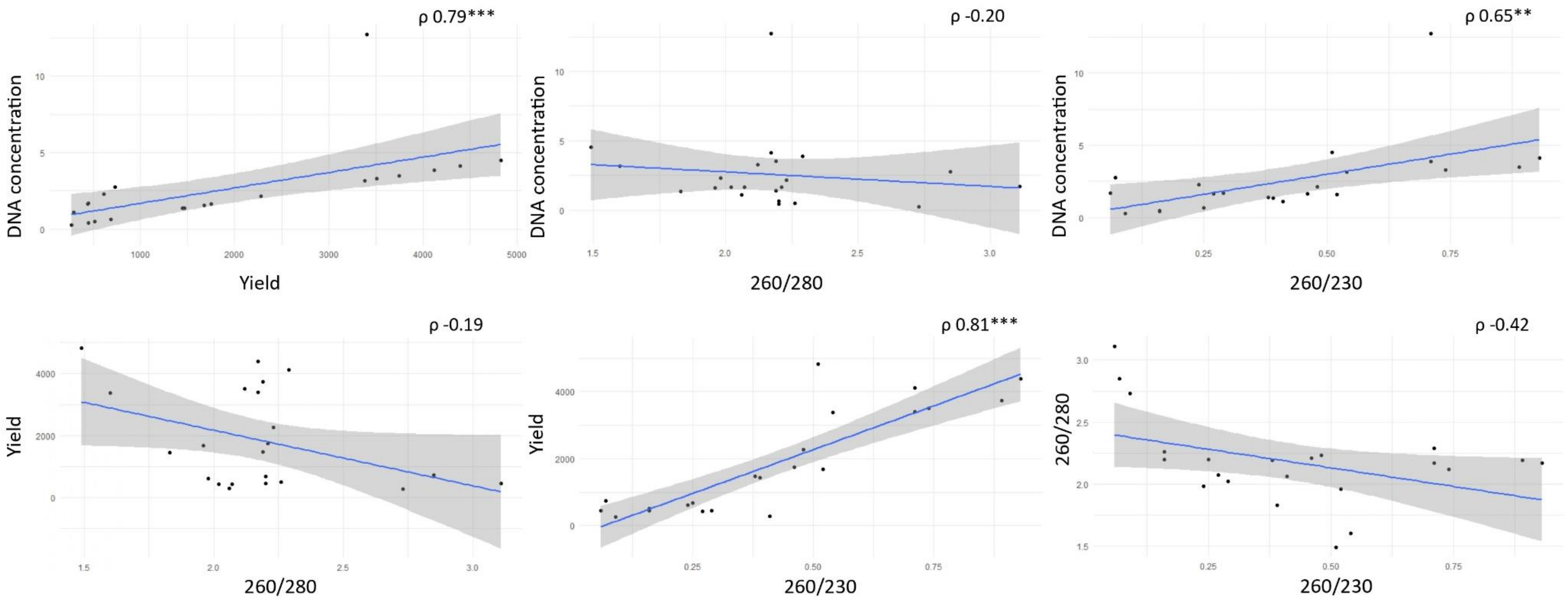


Figure S4. Spearman correlations between all pairs of extraction success variables measured for *Orthotomicus erosus*. Asterisks denote significant differences (*p < 0.05, **p < 0.01, ***p < 0.001)

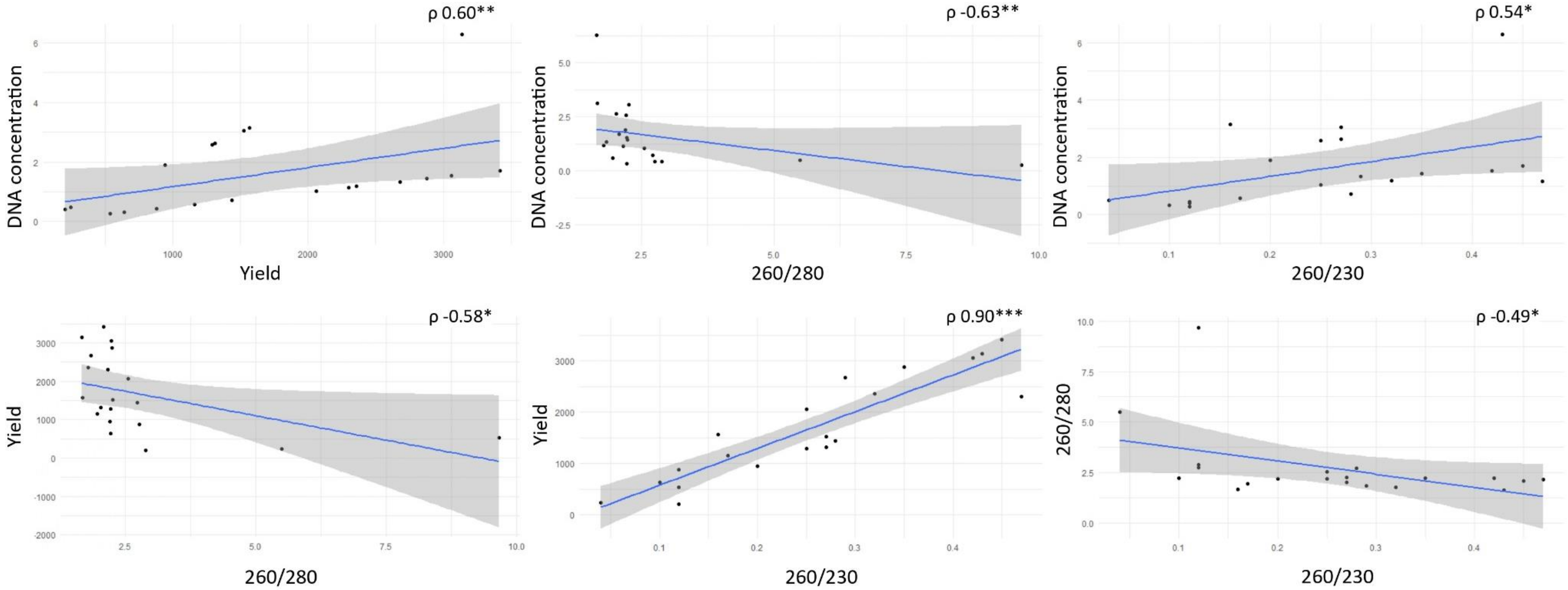


Figure S5. Spearman correlations between all pairs of extraction success variables variables measured for *Cyrtogenius luteus*. Asterisks denote significant differences (*p < 0.05, **p < 0.01, ***p < 0.001)
